# Glibenclamide, ATP and Metformin Increases the Expression of Human Bile Salt Export Pump ABCB11

**DOI:** 10.1101/2020.09.01.277434

**Authors:** Nisha Vats, Ravi Chandra Dubey, Madhusudana Girija Sanal, Pankaj Taneja, Senthil Kumar Venugopal

**Affiliations:** Institute of Liver and Biliary Sciences, D1 Vasant Kunj, New Delhi; South Asian University, Akbar Bhawan, Chanakyapuri, New Delhi; Sharda University, Plot No. 32, 34, Knowledge Park III, Greater Noida, UP

**Author notes:** **Corresponding Author:** Madhusudana Girija, Sanal MBBS, PhD Assistant Professor, Department of Molecular & Cellular Medicine, Institute of Liver and Biliary Sciences, D1 Vasant Kunj, New Delhi-110070.

**Keywords:** BSEP/ABCB11, ABCB11-KO, Insilico, upregulation, HepG2, glibenclamide, ATP, and metformin, Nuclear Receptors

## Abstract

**Background:** Bile Salt Export Pump (BSEP/ABCB11) is important in the maintenance of the enterohepatic circulation of bile acids and drugs. Drugs such as rifampicin, glibenclamide inhibit BSEP. Progressive Familial Intrahepatic Cholestasis Type-2, a lethal pediatric disease, some forms of intrahepatic cholestasis of pregnancy, and drug-induced cholestasis are associated with BSEP dysfunction.

**Methods:** We started with a bioinformatic approach to identify the relationship between ABCB11 and other proteins, microRNAs, and drugs. Microarray data set of the liver samples from ABCB11 knockout mice was analyzed by GEO2R tool. Differentially expressed gene pathway enrichment analysis was done by ClueGo v2.5.5 app from Cytoscape. Protein-protein interaction network was constructed by STRING application in Cytoscape. Networks were analyzed using the Cytoscape software v3.7.1. CyTargetLinker v4.1.0 was used to screen the transcription factors, microRNAs and drugs. Predicted drugs were validated on human liver cell line, HepG2. BSEP expression was quantified by Real Time PCR and Western Blot.

**Results:** ABCB11 knockout in mice was associated with a predominant upregulation and downregulation of genes associated with cellular component movement and sterol metabolism respectively. We further identified the hub genes in the network. Genes related to immune activity, cell signaling and fatty acid metabolism were dysregulated. We further identified drugs (glibenclamide and ATP) and a total of 14 microRNAs targeting the gene. Western Blot and Real Time PCR analysis confirmed the upregulation of BSEP on the treatment of HepG2 cells with glibenclamide, ATP, and metformin.

**Conclusion:** The differential expression of cell signaling genes and those related to immune activity in ABCB11 KO animals may be secondary to cell injury. We have found glibenclamide, ATP, and metformin upregulates BSEP. The mechanisms involved and the clinical relevance of these findings need to be investigated.

## INTRODUCTION

The Bile Salt Export Pump (BSEP), the major bile salt transporter in the liver canalicular membrane, is coded by *ABCB11* gene, and mutations in this gene cause Progressive Familial Intrahepatic Cholestasis Type-2 (PFIC-2) (1,2). Besides PFIC-2, mutations or insufficiency of BSEP is associated with a variety of diseases such as drug-induced cholestasis, pregnancy induced cholestasis, cryptogenic cholestasis, cholangiocarcinoma and hepatocellular carcinoma, which are cancers of the liver(3–7). Naturally, ABCB11 expression is induced by bile salts and is mediated by FXR-RXR heterodimer (8). Here in this pilot study we explored in-silico the interactions/networks around ABCB11. We wanted to identify the genes, drugs, microRNAs which might influence the expression of ABCB11. Drugs which could upregulate ABCB11 expression may be useful in ABCB11 haploinsufficiency and inhibition of the pump could result in the accumulation of toxic bile salts inside hepatocytes. Modulation of ABCB11 expression could be clinically beneficial in a variety of medical conditions.

## MATERIALS & METHODS

### Identification of differentially expressed genes

We analyzed the microarray data set of the liver samples from ABCB11 knockout mice (GSE 70179) using GEO2R online tool from NCBI (9). All differentially expressed genes (DEGs) were filtered with two criteria: −1> log_2_FC >+1 and adj. p-value <0.05.

### Pathway enrichment analysis

To identify DEGs which are significant, pathway enrichment analysis was conducted using the ClueGo v2.5.5 app from Cytoscape(10). ClueGo constructed and compared networks of functionally related GO terms with kappa statistics, which was adjusted at >0.4 in this study.

### Identification of hub genes and subnetwork analysis

The protein-protein interaction (PPI) networks were built by the Search Tool for the Retrieval of Interacting Genes (STRING v11.0) (11)and Cytoscape v3.7.1 software. The Molecular Complex Detection (MCODE), app from Cytoscape was used to screen modules of the PPI network with degree cut-off = 2, node score cut-off = 0.2, k-core = 2, and maximum depth = 100. The hub genes were identified by the CytoHubba app. The top 10 nodes were considered as notable hub genes and displayed.

### Identification of transcription factor and drug target

CyTargetLinker app from Cytoscape was used to identify the transcription factor (TF) and drugs targeting the dysregulated genes of the selected category. We drew Homo sapiens TF-target interactions linkset from database (ENCODE) (12) and drug-target interactions linkset from the database (DrugBank)(13). The networks were visualized and analyzed using the Cytoscape software v3.7.1

### Cell Culture

HepG2 cells were grown in high glucose DMEM (Hi-Media Lab, Mumbai, Cat. # AL111-500ML) supplemented with 10% fetal bovine serum (CellClone, Genetix Biotech Asia, New Delhi, Cat.# CCS-500-SA-U), penicillin and streptomycin (Hi-Media, Mumbai Cat. # A018-5X100ML). When cells became 80% confluent, they were treated with glibenclamide (500ng/mL) (14), metformin (25 mg/L) (15) and ATP (1mM) for 48h. Cells were harvested after 48h cells were harvested for total protein and RNA.

### Western Blot

Total proteins from HepG2 cells were prepared and run on 10% SDS-PAGE and transferred to a PVDF membrane using a transfer apparatus following the standard protocols (Bio-Rad). After incubation with 5% nonfat milk in TBST (10 mM Tris, pH 8.0, 150 mM NaCl, 0.5% Tween 20) for 1h the membrane was washed once with TBST and incubated with antibodies against human ABCB11 (Affinity, Catalog #DF 9278) 1: 2000 dilution; human β-actin (Santa Cruz Cat.# SC4778), dilution 1:1000. The membrane was washed and incubated with a 1:5000 dilution of horseradish peroxidase-conjugated anti-rabbit (Santa Cruz Cat# SC-2004)/anti-mouse antibodies (Cat.#SC-2005) for 2 h. Blots were washed with TBST four times and developed with the ECL system (Bio-Rad, US Cat.#170-5060) according to the ‘manufacturer’s protocol.

### Real Time PCR

Total RNA was isolated using NucleoZOL (Takara Cat. No. 740404.200) following manufacturer’s instruction. cDNA was prepared from (deoxyribonuclease treated) total RNA using RevertAid Reverse Transcriptase (Thermo Cat. No. EP0441) following the ‘manufacturer’s instructions. Real Time PCR was done with unique oligonucleotide primers targeting ABCB11 gene using GoTaq^®^ qPCR Master Mix (Promega Cat. No. A6001) following ‘manufacturer’s instructions on a Veriti Thermo Cycler from Applied Biosystems Waltham, Massachusetts, USA and data was acquired using the software platform associated with the same machine.

## RESULTS

### Expression of 560 genes changed significantly following ABCB11-KO in 1.5 m mice

Gene expression profile ABCB11 knockdown dataset GSE 70179 from GEO datasets were analysed with GEO2R tool. Genes with >2-fold change in expression value and <0.05 adjusted p value was filtered. Identified differentially expressed genes (DEG) from the GSE dataset were classified in two groups – upregulated (375 genes) and downregulated (185 genes) (Supp.Table-1). Gene ontology analysis was performed for functional analysis of DEGs by using ClueGo app from Cytoscape. PPIs of DEGs were constructed using STRING database showed an upregulation of genes related to cellular transport (pink colored nodes), and these nodes were also shared by Toll Like Receptor (TLR) signalling (Fig-1). Downregulated genes were involved in metabolic pathways (sterol, carbohydrate, alcohol, etc) (Supp. Table-2). We next identified top hub genes in PPI network using CytoHubba app from Cytoscape (Table-1). Immunologically important genes were among the top ranked upregulated hub genes (Fig-2a) downregulated group majorly represents cell signaling and fatty acid metabolism (Fig-2b). Epidermal Growth Factor Receptor (EGFR) ranked first among the genes involved in signaling pathways. Kinases play a role in the transcription, activity, or intracellular localization of ABC transporters as do protein interactions(16). Proteins which are interacting with ABCB11 are represented in Supp. Fig-1 which includes nuclear receptors NR1H4 and NR0B2. Most proteins were associated with bile acid metabolism and transport.

**Figure-1.**
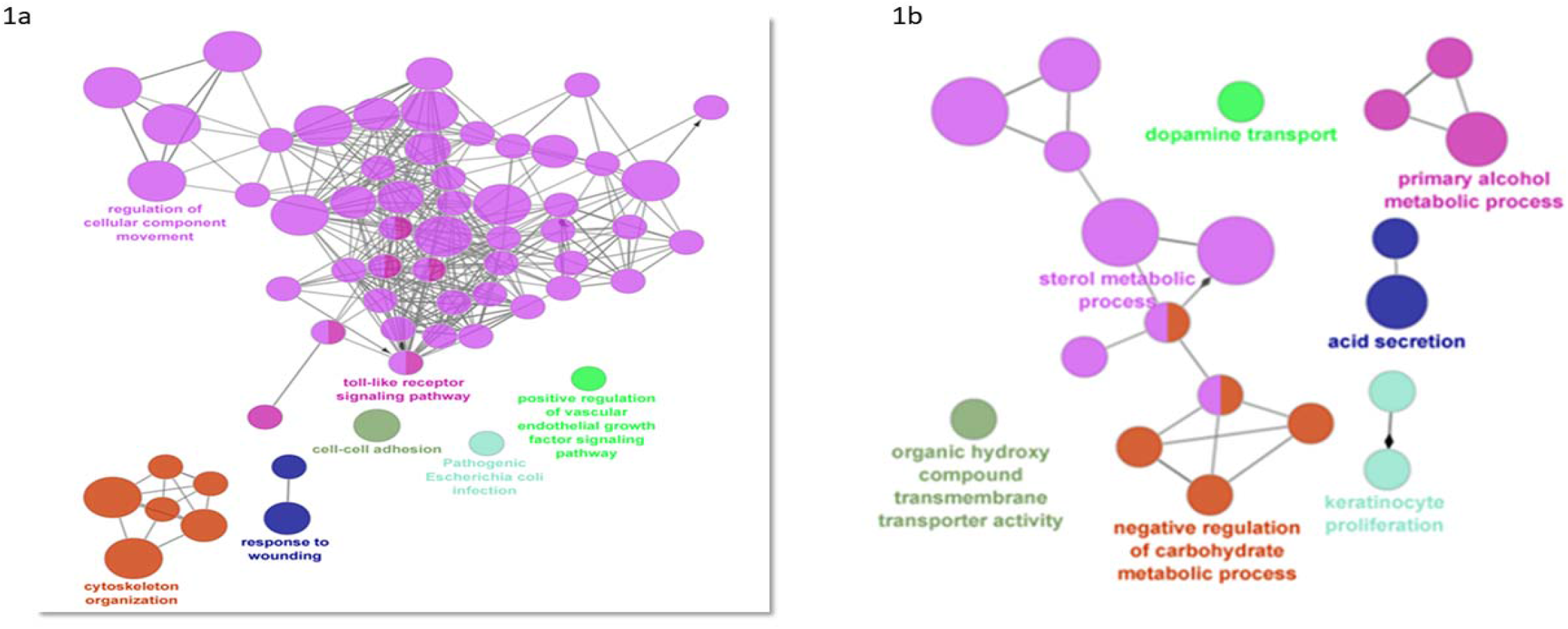
Protein-protein interaction networks (PPIs) of DEGs were constructed using STRING database. Gene ontology analysis was performed for functional analysis of DEGs by using ClueGo app from Cytoscape. This app allows simultaneous analysis of multiple annotation and ontology sources. Functionally grouped network is represented Figure 1a (upregulated genes) 1b (downregulated genes). The node size represents enrichment significance and connections are based on kappa score (> 0.4). In upregulated gene group maximum number of nodes which are in pink color represent the cellular component movement. These nodes are shared by toll like receptor signaling pathway.

**Table-1.**
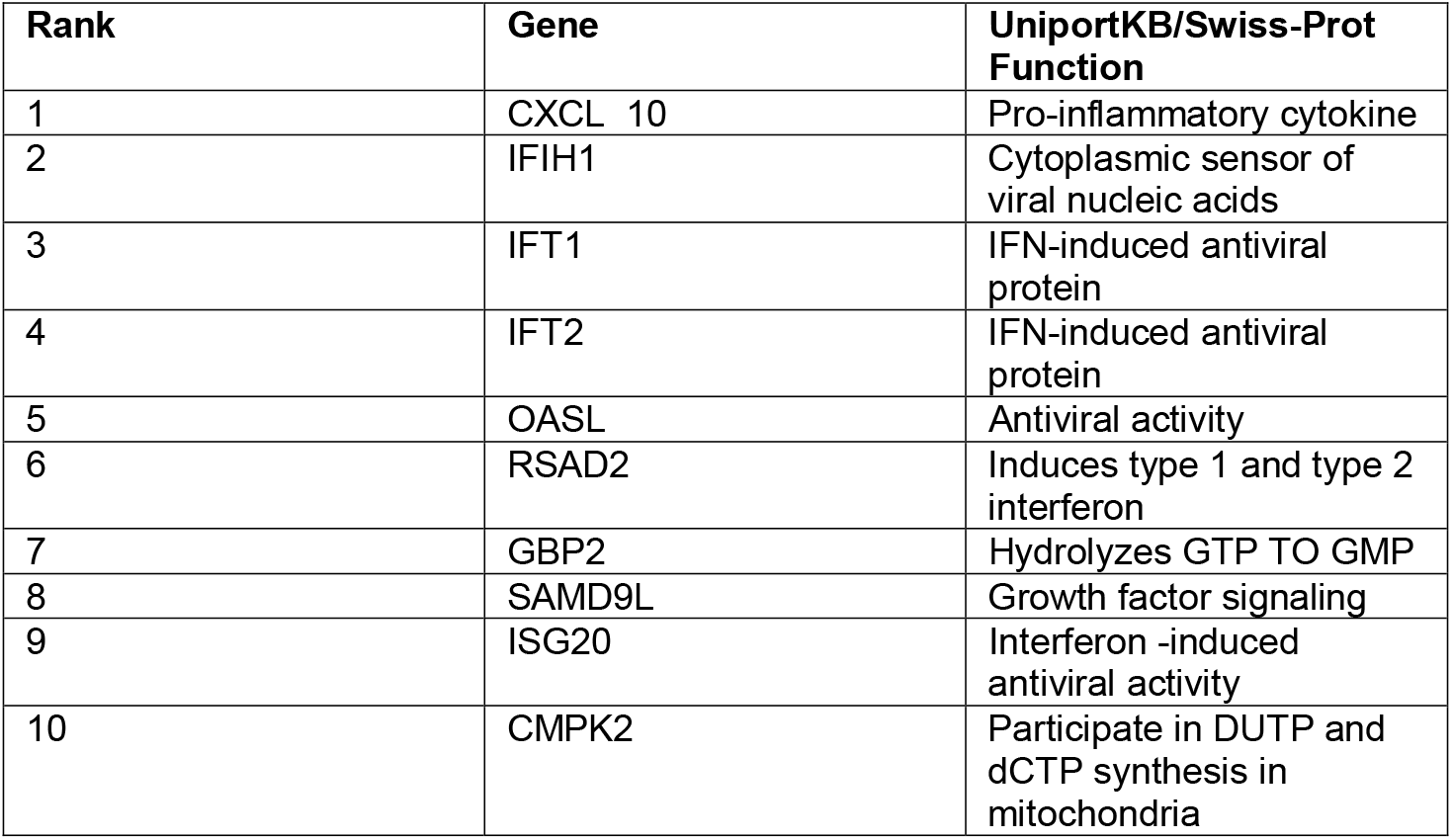
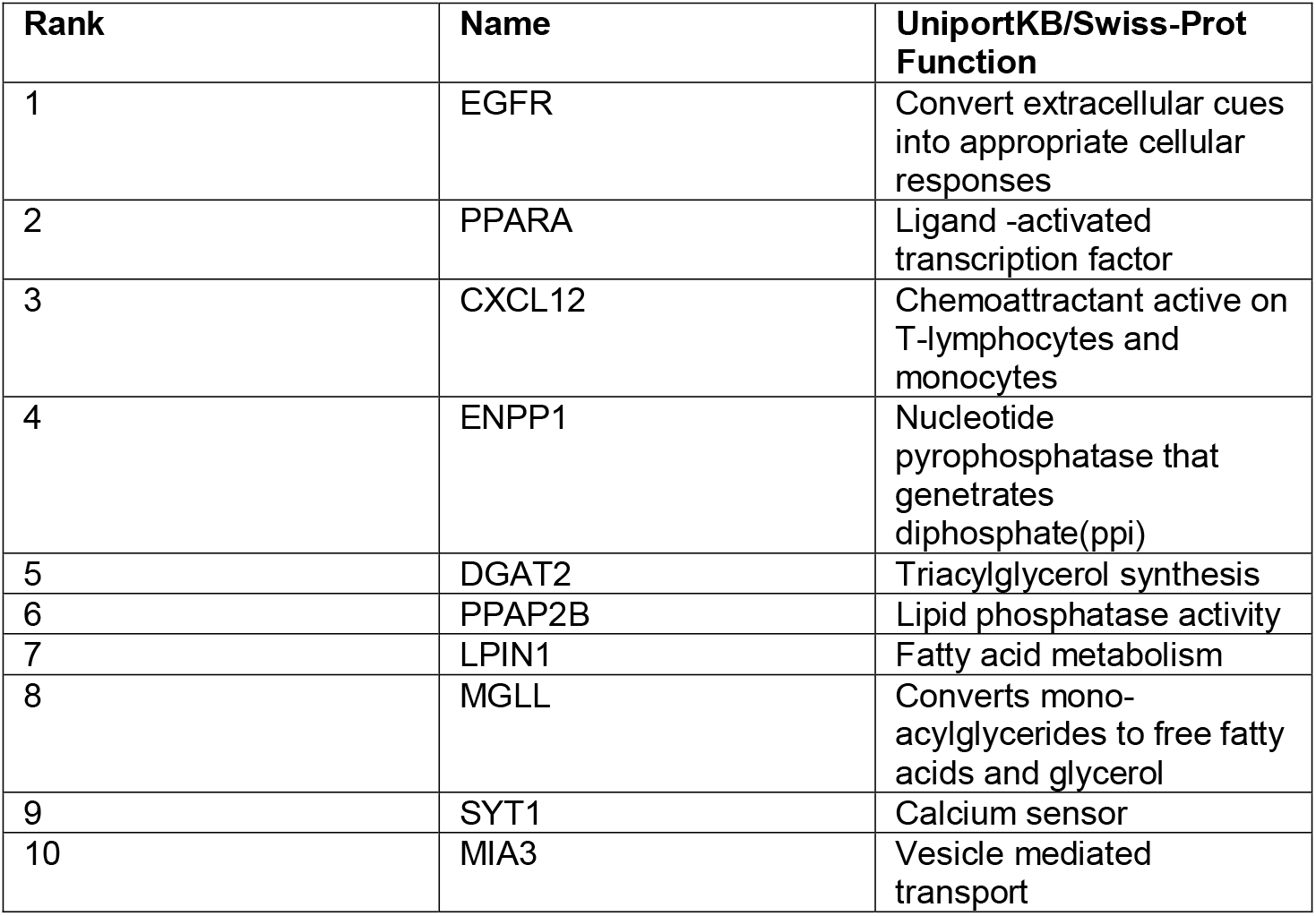
Table-1a Upregulated genes, Table-1b Downregulated genes. We observed that the top ranked hub genes in PPI network which were upregulated were associated with immune activity while those downregulated are associated with cell signaling and fatty acid metabolism. EGFR came first in the ranking which is a critical receptor in several cell signaling pathways.

**Figure-2.**
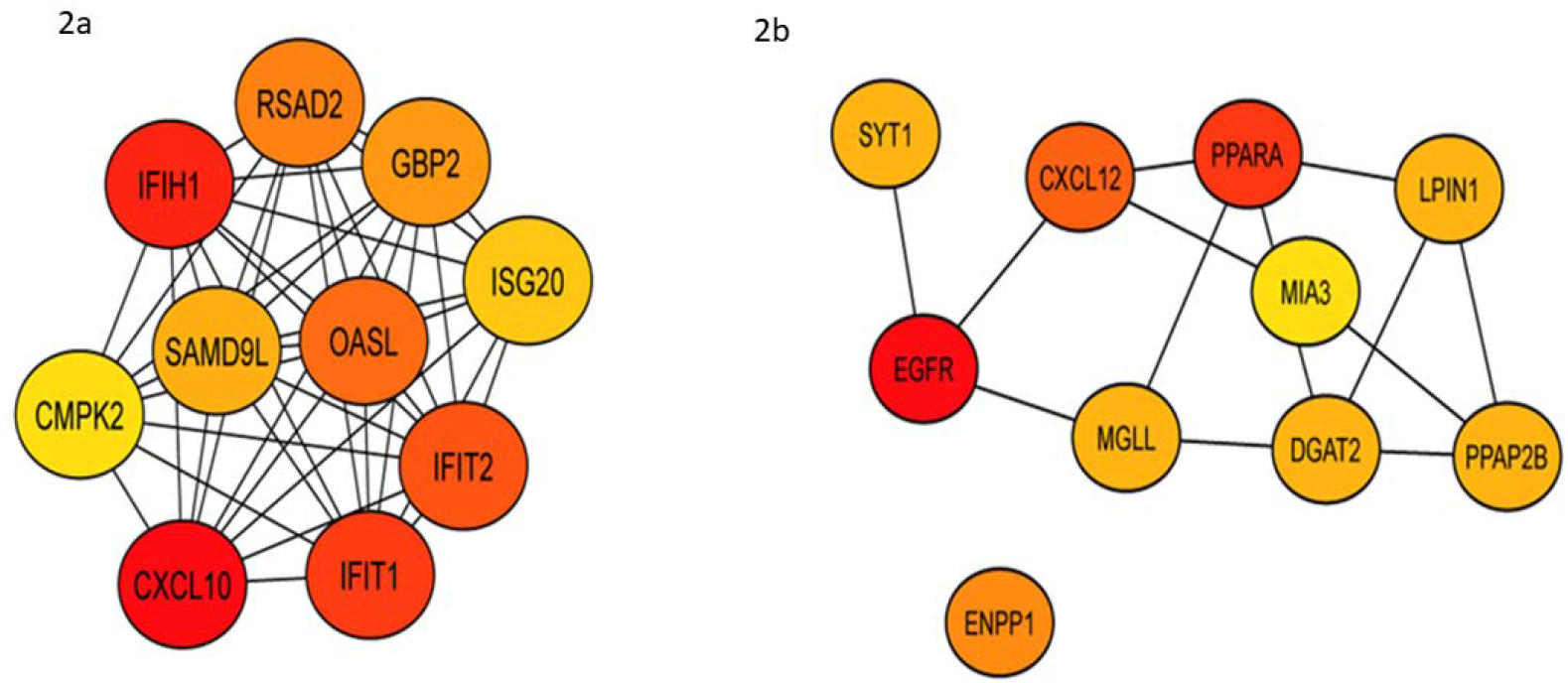
We identified top hub genes in PPI network of both upregulated and downregulated genes by using CytoHubba app from Cytoscape. We observed that the top ranked hub genes in the upregulated group were mainly related to immune activity (2a). The top hub genes in downregulated group were associated with cell signaling and fatty acid metabolism. The EGFR emerged as the top hub gene, a growth factor receptor which is crucial factor several cell signaling pathways (2b).

Sub-network analysis was performed using the Molecular Complex Detection (MCODE) app from Cytoscape (Fig-3), and CMPK2, ACTG1, and SSTR2 emerged as seed nodes among upregulated (Table -2). Among downregulated gene groups only one subnetwork was found to be significant which had three genes *MIA3* (which codes a protein which is important in the transport of cargos that are too large to fit into COPII-coated vesicles such as collagen VII), IGFBP4 (a protein that binds to both insulin-like growth factors and modifies their functions) and NOTUM (a carboxylesterase that acts as a key negative regulator of the Wnt signaling pathway by specifically mediating depalmitoleoylation of WNT proteins).

**Figure-3.**
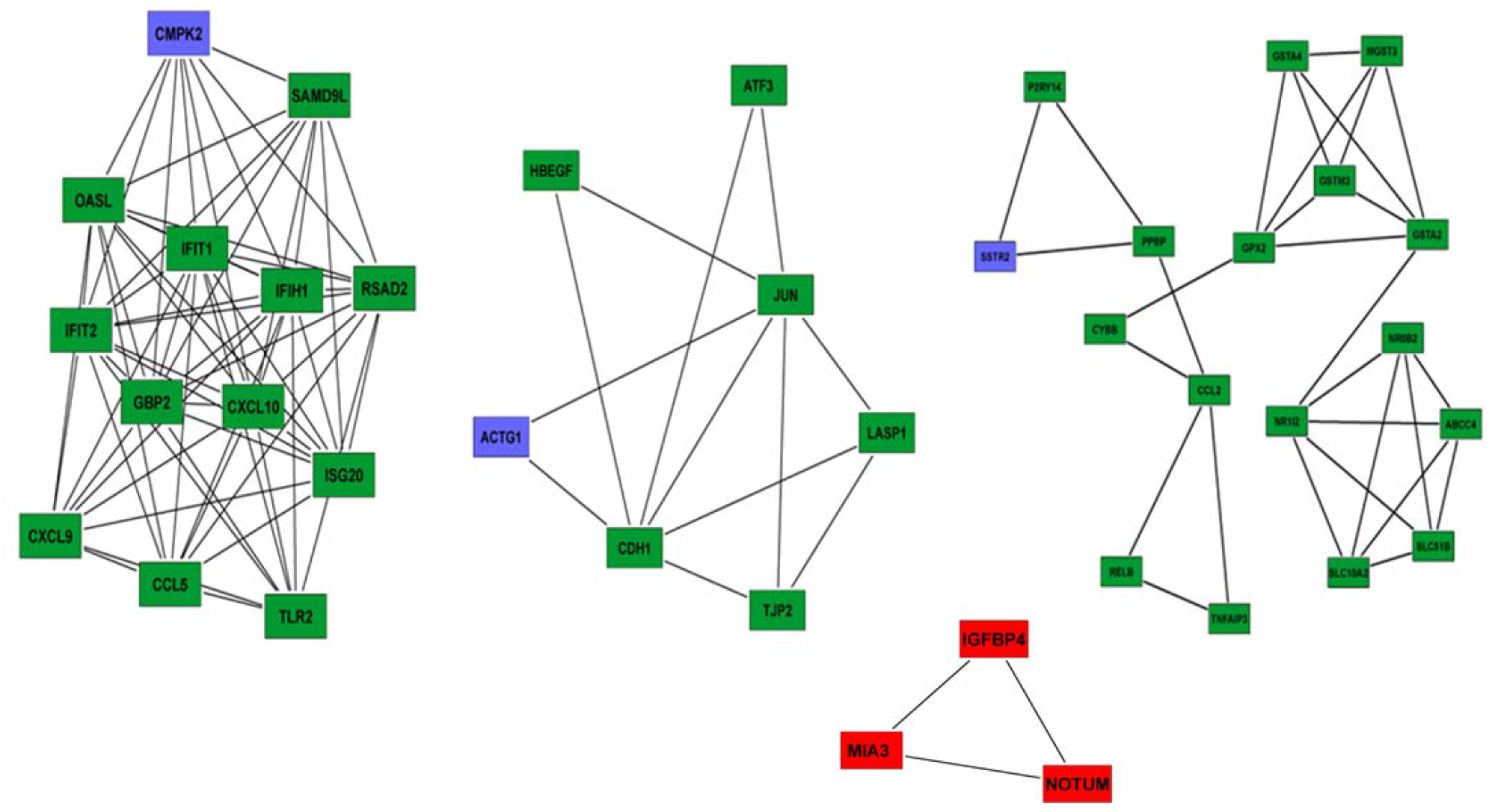
Sub-network analysis was conducted by using the Molecular Complex Detection (MCODE) app from Cytoscape. Top sub-networks on the basis of MCODE score (Degree cut-off= 2, node score cut-off = 0.2, k-core = 2 and max. depth = 100). Upregulated gene group clusters, we identified seed nodes (CMPK2, ACTG1 and SSTR2) in the network (green and blue). In downregulated gene group, we identified only one subnetwork which qualified cut off criteria. Three genes in this sub-network was identified: MIA3, IGFBP4 and NOTUM (red).

**Table-2.** Sub-network analysis. Sub-network analysis was performed using the Molecular Complex Detection (MCODE) app from Cytoscape and CMPK2, ACTG1 and SSTR2 emerged as seed nodes among upregulated

| **Sub-network** | **Seed-gene** | **UniportKB/Swiss-Prot Function** | **Diseases involved** |
| --- | --- | --- | --- |
| 1 | CMPK2 | This gene encodes one of the enzymes in the nucleotide synthesis salvage pathway that may participate in terminal differentiation of monocytic cells<br>May participate in dUTP and dCTP synthesis in mitochondria | Retinitis Pigmentosa 39 and Thiamine-Responsive Megaloblastic Anemia Syndrome |
| 2 | ACTG1 | Actins are highly conserved proteins that are involved in various types of cell motility | Deafness, Autosomal Dominant 20 and Baraitser-Winter Syndrome 2 |
| 3 | SSTR2 | Receptor for somatostatin-14 and -28 | Neuroendocrine Tumor and Growth Hormone Secreting Pituitary Adenoma<br>Cirrhotic liver and HCC express SSTRs. |

Using CyTargetLinker app from Cytoscape identified two drugs, glibenclamide, and ATP, directly targeting ABCB11. We subsequently looked for microRNAs [Target-scan database](17) that were associated with ABCB11, and a total of 14 microRNAs were identified targeting the gene (Fig-4). Transcription factors and microRNAs targeting ABCB11 and interacting partners are represented in Supp. Fig-2.

**Figure-4.**
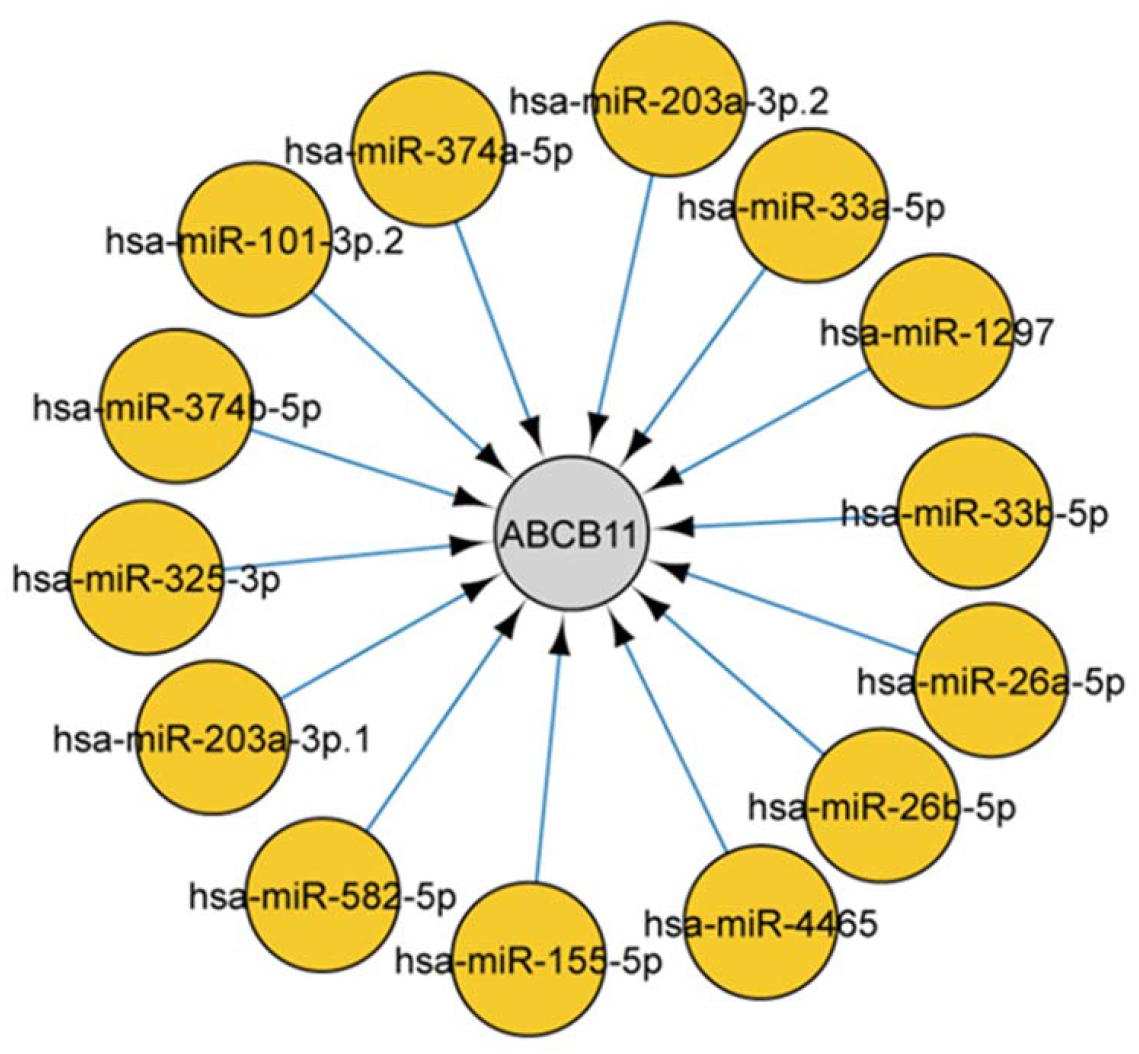
Cytoscape app CyTargetlinker version 4.1.0[6] was used to screen microRNAs using the Target-scan database. We identified microRNAs that were associated with ABCB11. In total 14 microRNA identified targeting the gene.

### Glibenclamide ATP and Metformin upregulates ABCB11

We evaluated in-vitro, the effect of three drugs, two of which were bioinformatically predicted (Glibenclamide, ATP) and one based on literature (18). We found all the three compounds upregulating ABCB11 expression based on Real Time PCR, and this was confirmed by Western Blot (Fig-5).

**Figure-5.**
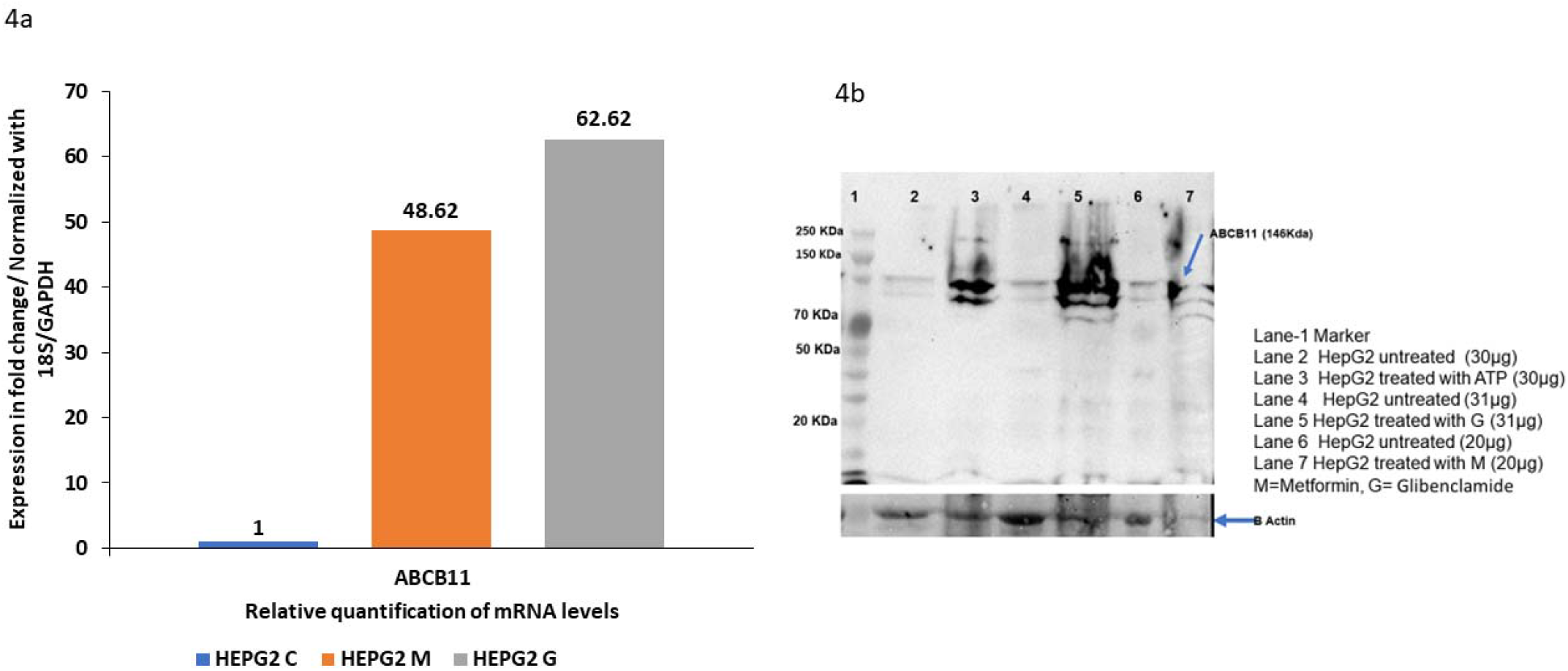
Three drugs, two of which were bioinformatically predicted (Glibenclamide, ATP) and one (Metformin) based on evidence from literature. All the three compounds upregulated ABCB11 expression based on Real Time PCR data. This was further confirmed by Western Blot against anti-human ABCB11 antibody as the primary antibody.

## DISCUSSION

We identified several immunologically important genes being upregulated ABCB11 deficiency. The reason could be liver cell injury secondary to bile salt accumulation, which triggers sterile immune response(19,20) and the downregulation of transport protein and metabolically important genes could be because of decreased liver function following damage. A regenerative response follows cell injury, and a host of genes involved in regeneration are upregulated (21–23) however, it appears that bile salts in the absence of BSEP hamper the regenerative response reflected by dysregulated collagen transporting protein MIA3 and NOTUM a protein involved in Wnt signaling. it’s also possible that EGFR is dysregulated via accumulating bile salts mediated by STAT3(24). We have observed an upregulation of ABCB11 in liver cell line (HepG2) on treatment with glibenclamide, metformin, and ATP. Its upregulation may be a compensatory mechanism in the case of glibenclamide and metformin because these drugs are known to inhibit ABCB11(25). Metformin is known to interfere with ABCB11 function mediated through AMPK-FXR crosstalk(18) involving metformin induced FXR phosphorylation. ATP acts through ATP receptors on hepatocytes(26,27). ATP is known to cross plasma membrane(28) and this can act via AMPK. However, ATP has a very short half-life (29), and it may be converted to ADP, which can activate AMPK(30). In a recent report, metformin was shown to suppress ABCB11 expression, which is not in agreement with our observation, however, they performed their experiment on primary human hepatocytes, and they have also treated their cells with DMSO (31). However, we are still unable to explain the observed difference. In conclusion, we need more experiments to determine the mechanisms of action of these drugs on the upregulation of ABCB11. Many changes in gene expression following ABCB11 could be secondary to stress, immune and regenerative responses following hepatocyte injury in mice liver.

## Disclosures

### Ethical approval and Informed consent

No human patient or animal material/specimens were used in this work which requires an Institutional Ethics Committee/Institutional Review Board approval to the best of our knowledge.

## Acknowledgement

The corresponding author (Sanal MG) acknowledges the Science and Engineering Research Board (**SERB**), Government of India and the Department of Biotechnology (**DBT**), Government of India for a limited financial support for about 18 months.

---

**Supp-Fig-1 a,b.**
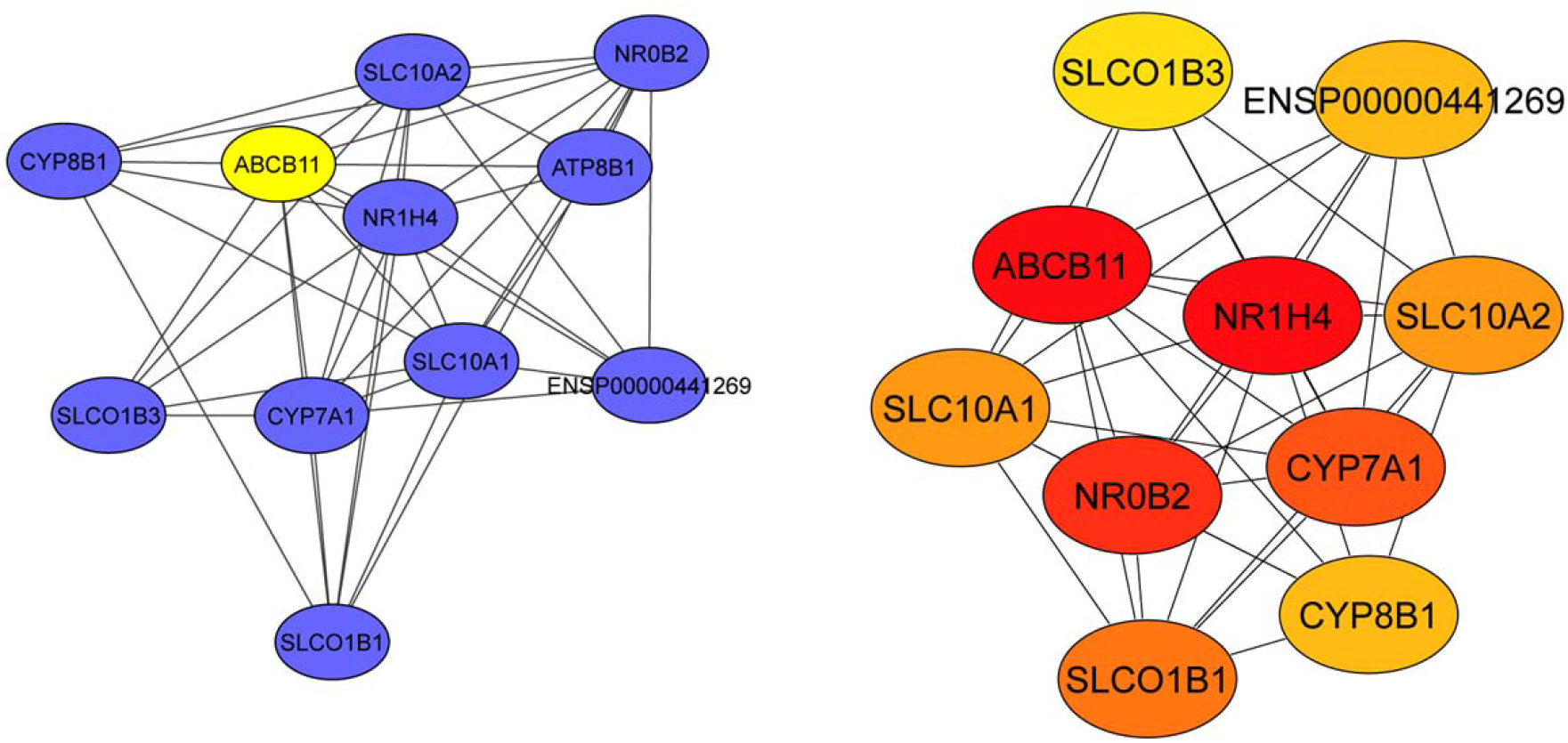
Protein-protein interaction network of ABCB11 gene was constructed by STRING app in Cytoscape with confidence score > 0.4 for homo sapiens. This network was constructed to analyse the relationship between ABCB11 and other proteins. Cytohubba app was used to calculate centrality of each node by MCC method. Node colour (red to yellow) represents the significance of the centrality in the group. In this analysis, we counted 11 nodes and 42 edges. These proteins majorly involved in bile acid metabolism and transport. Most of these genes are participant of more than one pathway which was expected because these pathways intersect and coregulated. We also mapped NR0B2 protein which is participate in sterol metabolism.

**Supp-Fig-2 a, b.**
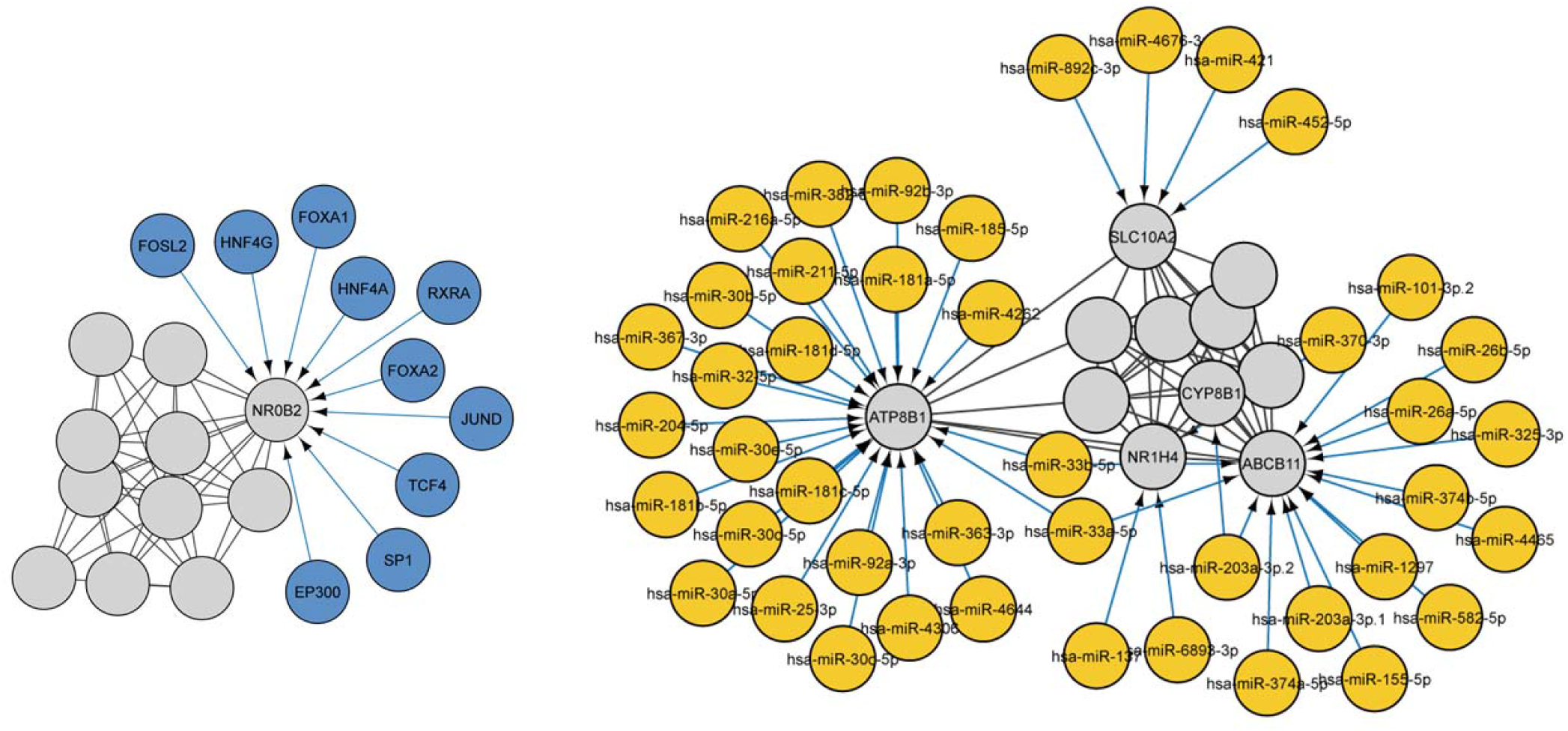
Cytoscape app CyTargetlinker version 4.1.0 was used to screen the transcription factors and microRNAs using ENCODE and Target-scan databases respectively. In the screen of transcription factors of ABCB11 interaction network we observed 21 nodes and 52 edges. Among these transcription factors, FOXA have been suggested an important factor in bile duct development and lipid accumulation. HNF4A in the regulation dyslipidaemia and terminal liver failure and JUND in fibrosis development. Others can be investigated in future studies. We counted 55 nodes and 89 edges in the search of microRNA targeting the ABCB11 network. Four genes (ABCB11, ATP8B1, SLC10A2 and NR1H4) targeted by multiple microRNAs also some microRNA such as has-miR-203a-3p.2 and has-miR-203a-3p.2 target more than one gene. By nature, a microRNA can regulate several pathways therefore it would be interesting to study in future the dysregulation of these microRNAs and interaction with Identified transcription factors.

